# CNAPE: A Machine Learning Method for Copy Number Alteration Prediction from Gene Expression

**DOI:** 10.1101/704486

**Authors:** Quanhua Mu, Jiguang Wang

## Abstract

Copy number alteration (CNA), the abnormal number of copies of genomic regions, plays a key role in cancer initiation and progression. Current high-throughput CNA detection methods, including DNA arrays and genomic sequencing, are relatively expensive and require DNA samples at a microgram level, which are not achievable in certain occasions such as clinical biopsies or single-cell genomes. Here we proposed an alternative method—CNAPE to computationally infer CNA using gene expression data. A prior knowledge-aided machine learning model was proposed, trained and tested on the transcriptomic profiles with matched CNA data of 9,740 cancers from The Cancer Genome Atlas. Using brain tumors as a proof-of-concept study, CNAPE achieved over 90% accuracy in the prediction of arm-level CNAs. Prediction performance for 12 gene-level CNAs (commonly altered genes in glioma) was also evaluated, and CNAPE achieved reasonable accuracy. CNAPE is developed as an easy-to-use tool at http://wang-lab.ust.hk/software/Software.html.

## 1 INTRODUCTION

COPY number alteration (CNA) represents the abnormal number of DNA segments. Considering the size of abnormalities, CNA can be divided into broad CNAs or aneuploidies, which stand for the aberrant number of chromosomes or chromosomal-arms, and focal CNAs which stand for the amplification or deletion of smaller genomic regions. With the development of next-generation sequencing technology, recent genomic landscape studies have revealed that CNAs exist in about 90% of all human cancers [1]. For instance, glioblastoma (GBM), the most common and aggressive brain cancer in adults, is characterized by the amplification of chromosome 7 and the deletion of chromosome 10 [2–3]. These alterations are found in about 66% of the GBM cases [4].

The earliest attempts to detect CNA in human cancer date back to over 100 years ago [5], when scientists observed the stained mitotic cells that were arrested in the metaphase or prometaphase of the cell cycle under the light microscope. Recent microarray-based methods, such as the single nucleotide polymorphism (SNP) array and the comparative genomic hybridization (CGH) array, were developed. The CGH based methods detect CNA by comparing the fluorescence ratio with the normal DNA [6], while the SNP-array based methods calculate the CNA by modeling the allele frequencies of a large number of common SNPs along the genome [7]. Both methods substantially improved resolution and accuracy. More recently, methods based on DNA sequencing (DNA-seq) data were developed, assuming that the number of sequences aligning to each genomic region is proportional to its copy number [8]. Yet this approach is complicated by GC content, the mappability of the genome, as well as sample purity [9–11]. Later development incorporating both read depth and allelic frequency significantly improved the method performance, even in low purity samples [12]. However, DNAseq data are not always available, especially in clinical applications where the total number of cells in biopsies are limited.

Computational inference of CNA from RNA sequencing (RNA-seq) provides an alternative solution. Compared to DNA-seq, RNA-seq usually requires less biological materials, has a relatively lower price, and more importantly, it provides additional information including not only transcriptomics but also genomics and microenvironments of targeted samples, thus is more desirable for a wide range of studies. As a result, more RNA-seq data are available than DNA-seq in public repositories, such as NCBI Short Read Archive. Especially for single-cell sequencing studies, RNA-seq has been widely applied while simultaneously sequencing DNA and RNA from an individual cell remains immature [13–14].

However, the inference of CNA from RNA-seq is challenging given that the regulatory machinery of different genes highly varies. This is further complicated by a number of regulatory mechanisms, such as the activity of transcription factors, the status of DNA methylation, chromosomal openness and so on. Computational technologies similar to that used in the previously mentioned SNP-based CNA detection method could be applied, but the SNPs in certain regions, especially the deleted ones, might be overlooked due to their low expression. In 2014, Patel *et al* proposed a sliding-window smoothing method to infer CNA from single-cell RNA-seq [15]. This method sorted all genes by their genomic coordinates, followed by the normalization of gene expression in each cell against normal control, and then computed the median of the normalized expression values in every 50-gene window as the indicator of DNA copy numbers. This simple method has been applied in several subsequent studies [16–19]. Hu *et al* adopted this method to predict 1p/19q co-deletion from bulk RNA-seq data of gliomas and achieved about 95% accuracy [20]. However, this method assumes the impact of the CNA events to be limited to the corresponding amplified/deleted regions, which is usually not the case, especially when the copy number of regulators or transcription factors were altered. Moreover, the window size is fixed and artificial, which might limit its scope of application. Wang et al [21] compared five existing methods for predicting 1p/19q co-deletion in gliomas, and they concluded that smoother method was the most stable approach, which achieved high accuracy (97.8% and 98.6%) in two independent cohorts. Yet this study did not investigate the prediction of other CNAs.

In this work, we reanalyzed RNA-seq data from over 9,000 human cancer samples labeled by matched CNA information and trained prior-knowledge aided machine-learning models to predict both large-scale and single-gene CNAs from gene expression. In Section 2 we detail the motivation and mathematical formulation of our method, including the details of model training and testing; in Section 3 we present the results of both large-scale and single-gene CNA prediction models. In Section 4 we summarize and raise potential future development of our method.

## 2 MATERIALS AND METHODS

### 2.1 Correlation between CNA and gene expression

Gene expression is a complicated process under the deliberated regulation of many factors, such as transcription factor activity, DNA methylation, chromatin status, and copy number. To explore the impact of CNA on gene expression, we compared brain tumor samples with 1p/19q co-deletion (N=169) and those without such co-deletion (N=491) from the TCGA pan-glioma cohort [22]. We observed that in the samples with 1p/19q co-deletion, while 1462 genes were significantly down-regulated (log_2_(fold change) < −1 and FDR-adjusted *q*-value<0.05), 879 genes were significantly up-regulated (log_2_(fold change) > 1 and FDR-adjusted *q*-value<0.05, Fig.1a). Among 2341 significantly differentially expressed genes, 289 located on chromosome 1p or chromosome 19q. Surprisingly, we observed 11 genes on chromosome 1p or chromosome 19q were significantly up-regulated (to the opposite direction of the copy number change) with log2(fold change) > 1.5. These genes were *TNNT1* (19q), *PTGFR* (1p), *EPHA10* (*1p*), *NT5C1A* (*1p*), *HTR6* (1p), *SLC8A2* (19q), *FXYD7* (19q), *ASPDH* (19q), *SLC6A17* (1p), *GABRD* (1p) and *SHISA7* (19q). Gene ontology (GO) analysis showed significant enrichment in “integral component of plasma membrane” (GO: 0005887, P=2.7×10^−4^, hypergeometric test).

**Fig. 1.**
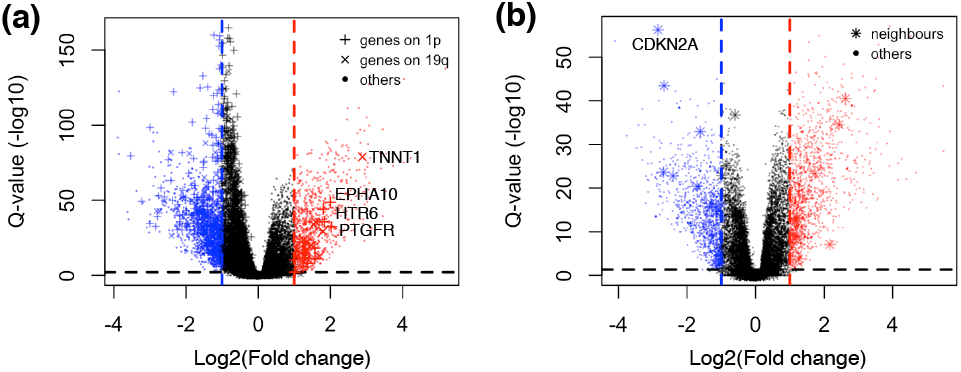
Relationship between gene expression and copy number alteration. (**a**) Volcano plot showing the differential expression genes (DEGs) in 1p/19q co-deleted gliomas compared to non-codel samples. The dashed lines show the significance threshold: absolute value of log2(fold change) > 1 and FDR-adjusted q-value < 0.05. (**b**) DEGs in gliomas with *CDKN2A* deep deletion compared to those without *CDKN2A* deletion. Genes located within 20 Mb of *CDKN2A* were defined as its neighbors.

In addition, we performed a similar analysis for a genelevel alteration, *CDKN2A* deep deletion in the same cohort. In total 136/660 samples have *CDKN2A* deletion. We found 1291 significantly up-regulated genes and 995 down-regulated genes. Defining genes with distance to *CDKN2A* less than 20 Mb as its neighbors, we found 16 of the 170 genes were significantly down-regualted, while 9 were significantly up-regulated (Fig. 1b).

The differential gene expression demonstrated that (i) the impact of CNA is not limited to the deleted/amplified regions; (ii) the changes in gene expression do not always follow the direction of CNA. To disentangle such complex correlation between the change of gene expression and CNA, a more comprehensive method is necessary to be developed.

### 2.2 Machine-learning model for CNA prediction from gene expression data

On the basis of the above findings, we proposed CNAPE, a multinomial logistic regression model with least absolute shrinkage and selection operator (LASSO) and knowledge-based feature selection to infer CNA from gene expression data [23]. Given a genomic region, the possible copy number status of this region could be normal (N), amplified (A), or deleted (D). Define the expression data of *p* genes in *n* samples as *X*_*p*×*n*_ and the corresponding copy number status of one genomic region *R* in these samples as *Y*_1×*n*_. The CNA status in this genomic region is modeled as follows

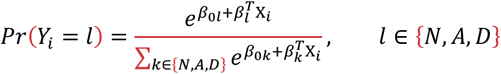

Here X_*i*_ is the gene expression profile of sample *i*, *l* corresponds to the CNA status, and *β*_*l*_ is a vector containing the weight of the *p* genes corresponding to status *l*. To select biologically meaningful and most informative genes, a weighted penalty on the scale of *β*_*p*×*k*_ is applied. In particular, the loss function to optimize *β* for region *R* is defined as

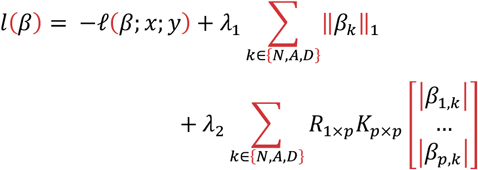

here *ℓ* is the maximum likelihood function, λ_1_ is the *L*_1_ norm hyper-parameter, determined during cross-validation training; *λ*_2_ is a hyper-parameter that indicates the confidence level on biology knowledge; *R*_1×*p*_ is an indicator of whether the candidate genes locate at the targeted region *R*; *K*_*p*×*p*_ indicates interactions between genes defined by biological prior knowledge. We set *λ*_2_ to 0 for chromosomeand arm-level CNA prediction models because our knowledge about the function of these alterations is minimal. For predicting gene-level CNA of a specific gene, protein-protein interaction networks as annotated in the STRING database are extracted and inputted as *K*_*p*×*p*_. For *R*_1×*p*_, we considered various numbers of nearest genes, and the optimal number is selected via cross-validation. The optimization process is carried out in R using the “glmnet” package [23].

### 2.3 Model training and testing

We acquired the cancer genome atlas (TCGA) dataset from the PancanAtlas publication page (https://gdc.cancer.gov/about-data/publications/pan-canatlas), in which for 9,740 samples from 33 cancer types both gene expression and CNA profiles are available [1]. In particular, the copy number of each autosome and each chromosome arm (except the short arms of chromosome 13, 14, 15, 21 and 22 since they contain less than 50 genes) in each sample is annotated as either amplified, deleted, neutral, or not applicable (NA). Here, NA means that a single value is not appropriate to summarize the complex pattern of one chromosome where amplifications and deletions co-exist.

A nested cross-validation strategy was used to train and test the models. The data were randomly divided into three parts, two of which were used for training and the third part for testing. In the training process, five-fold cross validation was employed. The trained model was then applied to the third part of the data to evaluate the performance. To test robustness, the training and testing process were performed iteratively for three to six times by randomly reassigning training and testing cohorts, and the results were reported as follows.

## 3 RESULTS

### 3.1 Chromosome-level models

Accuracy, the number of correct predictions divided by the total number of predictions, was used as the evaluation metric. The performance of the chromosome level models was summarized in Fig. 2a. Overall, the accuracy on the testing dataset was higher than 80% for each chromosomal CNA. For chromosomes 8 and 13, the median accuracy is higher than 90%. The accuracy was higher than 85% for 19 chromosomes. The lowest accuracy was observed in chromosome 1 (median: 81.5%), possibly due to the large size of this chromosome.

**Fig. 2.**
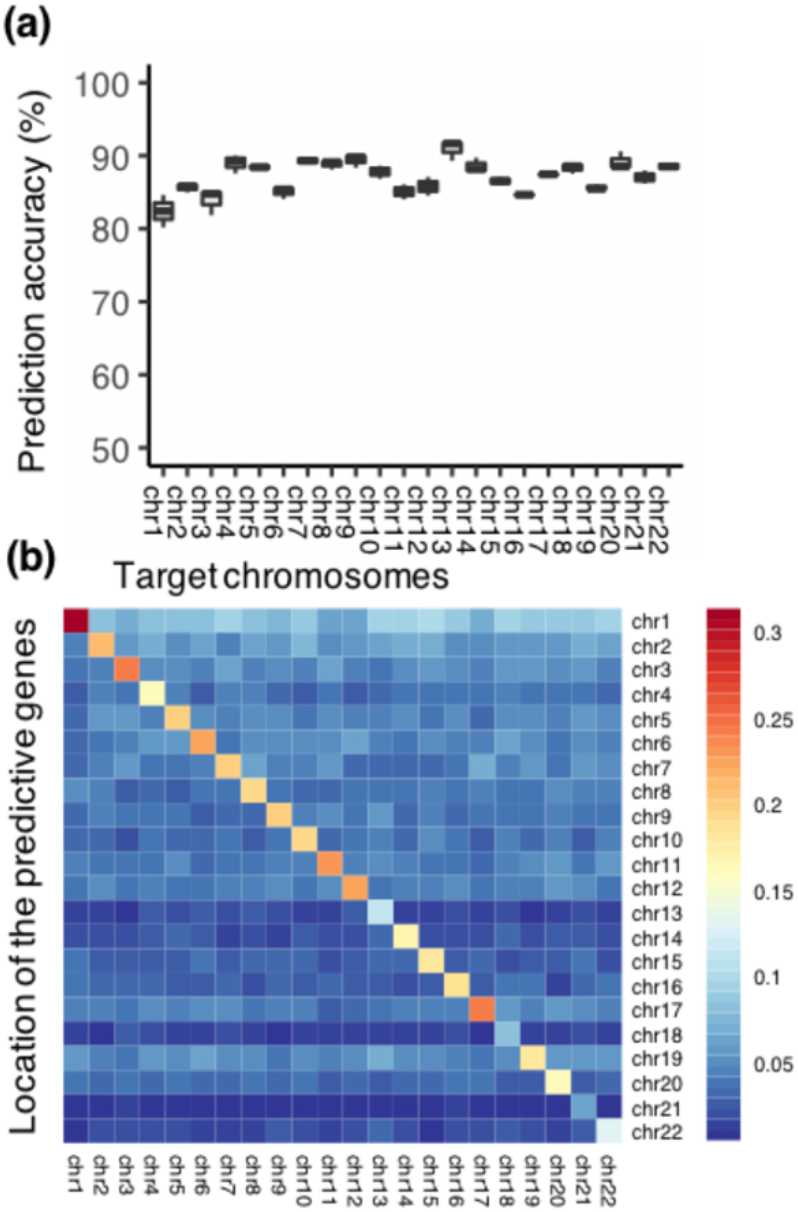
(**a**) Prediction accuracy of the chromosome-level models in TCGA dataset. (**b**) Distribution of the feature genes for each chromosome-level model. Each column stands for one prediction model, and for each model, the rows stand for the locations of the feature genes. Each cell represents the proportion of feature genes on corresponding chromosome.

In all chromosomal copy number prediction models, we analyzed the feature genes, defined as genes for which |*β*| > 0 in the model, and found that the feature genes were highly enriched in the target chromosome (Fig. 2b). For instance, in the model predicting CNA of chromosome 1, 31.4% of the feature genes locate on chromosome 1. However, there are also feature genes locate on other chromosomes, suggesting the non-local genes should be also considered in the CNA prediction.

### 3.2 Chromosome arm-level models

The performance of the arm-level models was summarized in Fig. 3a. Independent testing showed that arm-level prediction models achieve an average accuracy 88.4% (± 2.0%). Not that accuracy of all arm-level models is higher than 85%, with >90% for eight chromosome arms. Compared to the models for chromosomal CNA, the arm-level models showed higher accuracy.

**Fig. 3.**
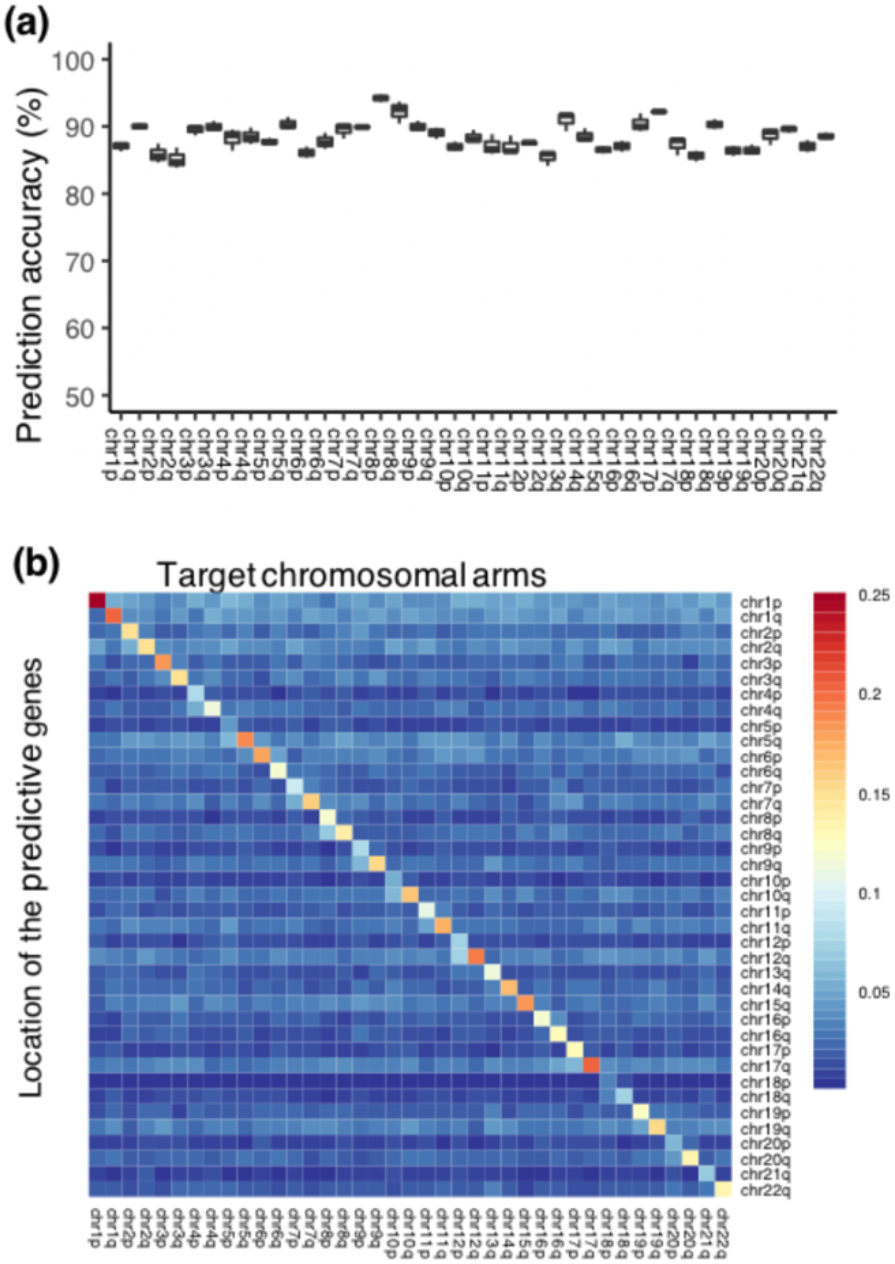
(**a**) Prediction accuracy of the arm-level models in TCGA dataset. (**b**) Distribution of the selected feature genes of each arm-level CNA prediction.

Similar to the observations in chromosome-level models, the feature genes in each model were enriched in the target arm or its neighbor arm. About 5.0% to 25.1% feature genes were located on the target arm, while some other feature genes locate outside the target chromosome (Fig. 3b).

### 3.3 1p/19q codel prediction

We then applied our model to predict a broad CNA, chromosomes 1p and 19q co-deletion (1p/19q codel). 1p/19q codel is one of the key molecular features for glioma diagnosis and subtyping. Glioma patients with this alteration show distinct cell morphology in histology, and they have significantly better prognosis under standard treatment [24–26].

The TCGA pan-glioma dataset included 660 samples for which both RNA-seq and 1p/19q codel status were available. In this cohort 169 (25%) of the cases were 1p/19q positive. Using the same framework, we trained the model on randomly selected 80% of all samples and tested on the remaining 20%. Our model obtained an AUC of 0.997 (sensitivity: 1, specificity: 0.996, Fig. 4a). We applied the model to the whole TCGA pan-glioma dataset and found three samples were mis-predicted. We therefore checked the details of the three gliomas (Fig. 4b). An alternative breakpoint was found in TCGA-CS-5394, where a segment of about 7Mb in 19q was retained. The strict definition of 1p/19q co-deletion is not quite clear, but our method clearly captured the copy number change in the major segments. The label of TCGA-VM-A8CA in the initial annotation was mistakenly labeled as non-codel. Finally, TCGATM-A84R was mis-predicted as non-codel probably due to the sub-clonal mutation or lower tumor purity as evidenced by the allele frequency of the known clonal *IDH2* R172 mutation (0.21).

**Fig. 4.**
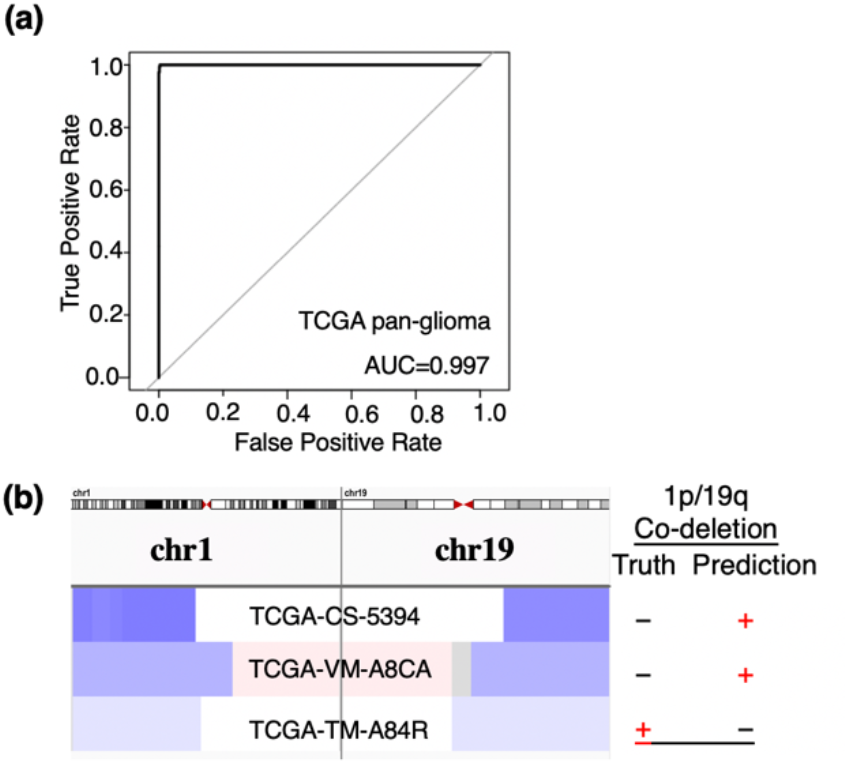
(**a**) Receiver operating characteristic (ROC) curves of the 1p/19q codel prediction model in TCGA pan-glioma testing cohort. The three different-color curves show three random partitions of training-testing runs. (**b**) Heatmap showing the copy number of chromosomes 1 and 19 in the three mis-predicted samples. Blue color represents deletion, and red color represents amplification. Color gradient indicates the extent of copy number changes.

### 3.4 Gene-level CNA inference

We further evaluated the performance of inferring genelevel CNA using our method. In particular, we selected 12 well-known glioma driver genes (*EGFR*, *MET*, *PDGFRA*, *CDK4*, *MDM2*, *CDKN2A*, *PTEN*, *TP53*, *CIC*, *FUBP1*, *RB1* and *NF1*) [27], and then trained and tested models to predict their CNAs in the TCGA pan-glioma dataset [22]. By adjusting *K*_*p*×*p*_ in the models, two types of candidate feature genes were considered: genes at nearby loci of the target gene, and genes functionally related to the target gene as defined in the STRING database.

One challenge in gene-level CNA prediction was that the positive/negative labels were imbalanced. Prevalent CNAs such as *PTEN* deletion was present in over 30% of patients, but some alterations such as *MDM2* amplification was observed in less than 10% cases. To evaluate the prediction performance in this dataset, AUPRC (Area Under the Precision-Recall Curve) is a more proper metric [28].

By adjusting *K*_*p*×*p*_ to control the candidate feature genes, we designed 10 groups of models, each considered 5, 10, 20, 50 or 100 neighbor genes, and with or without the functionally related genes, to compare and select the optimal combination of candidates. For each combination, six random training-testing partitions were performed. Taking *CDKN2A* deep deletion as an example, for models considering only neighbor genes as candidates, AUPRC was higher in models using 50 or 100 neighbor genes than models considering fewer genes (Fig. 5a). In addition, AUPRC were improved when functionally related genes were considered in addition to neighbor genes. To select the optimal combination of candidate feature genes, we calculated the average AUPRC among six randomizations, and models considering 50 neighbor genes plus functionally related genes as candidates showed highest mean AUPRC, therefore was selected as optimal. In the optimal model, 24 genes had non-zero weight, and ten of them were functionally related genes (Fig. 5b). The weight of each feature gene was shown in Fig. 5c. The gene with the most negative weight was *CDKN2A* itself, followed by *CDKN2B* and *MTAP* (close neighbors of *CDKN2A*), while the gene with the most positive weight was *CDK6* (located at 7q 21.2).

**Fig. 5.**
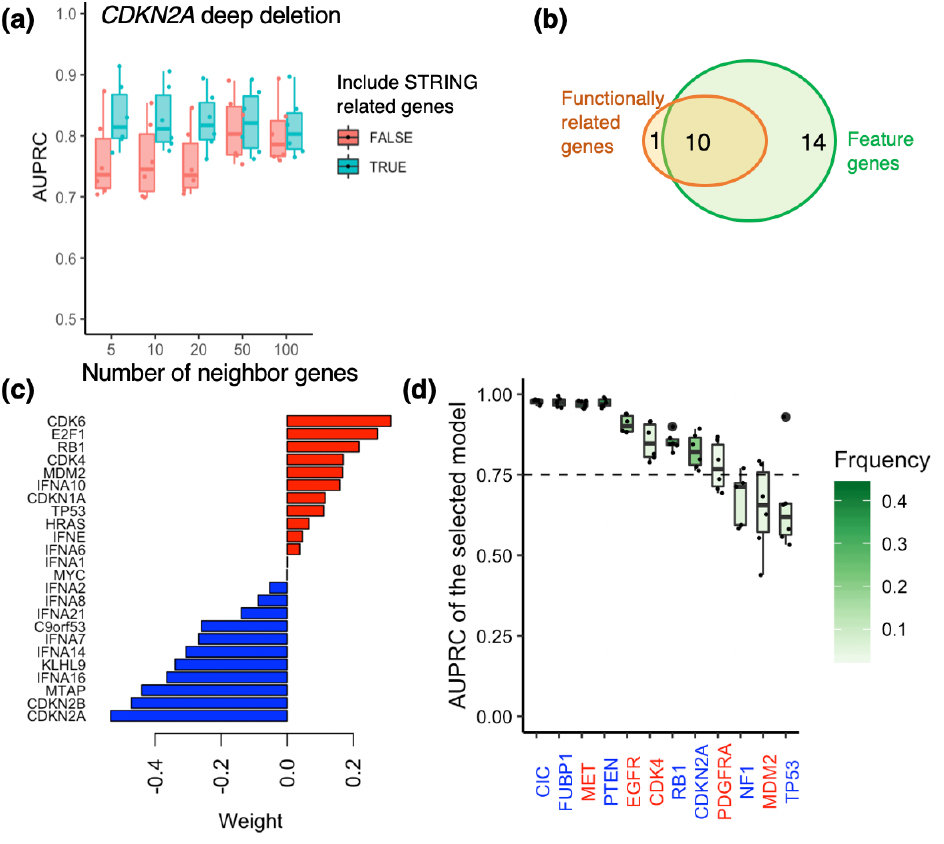
Single gene CNA prediction models. (a) Performance of *CDKN2A* deep deletion prediction models, considering various numbers of neighbor genes, or neighbor genes plus functionally related genes from the STRING database. Each dot represents one run with a random partition of training-testing data. The model with the highest mean AUPRC is selected as the best model and analyzed in b and c. (b) Overlap between feature genes in the selected model, and functionally related genes of *CDKN2A* in the STRING database. (c) Relative weight of feature genes in the selected model. (d) AUPRC of the 12 selected gene-level CNAs. For each gene, the blue color of the gene name indicates deletion, while red color indicated amplification. The horizontal line shows a cutoff at 0.75. A color gradient is used to show the frequency of each CNA.

The overall prediction accuracy varied from gene to gene (Fig. 5d). Among the 12 selected gene-levels CNAs, 8 (66.7%) had AUPRC > 0.75, including deletion of *CIC*, *FUBP1*, *PTEN*, *RB1* and *CDKN2A*, as well as amplification of *MET*, *EGFR* and *CDK4*. They also showed less variation compared to the other four CNAs, indicating that these models were more robust. Prediction AUPRC of *MDM2* amplification, PDGFRA amplification and *NF1* deletion were lower, probably due to their low incidence (6%, 10%, and 6%, respectively). *TP53* deletion had prediction AUPRC of 0.65 ± 0.15. In glioma, *TP53* loss of heterozygosity was known to be frequent, which may partly explain the lower average AUPRC and higher variation.

## 4 CONCLUSION AND DISCUSSION

While the idea of inferring CNA from RNA-seq data seems natural, this task is complicated by the variation in gene expression and regulatory mechanisms of transcription. Here we present a prior knowledge-aided machine learning model trained from the big data of TCGA for CNA inference in human cancer, using knowledge from protein/gene networks to guide feature selection. Our testing analyses demonstrated high accuracy in large-scale CNAs such as aneuploidy and arm-level alterations, which will be useful on occasions when DNA-seq data are not available. For single gene CNA, our study showed that the performance of this method varies from gene to gene. For the eight most common single gene CNAs in glioma, the AUPRC of our method achieved >0.75. However, the performance for the prediction of *TP53* deletion, *MDM2* amplification, *NF1* deletion and *PDGFRA* amplification, was lower than 0.75, and the models for other cancer-related CNAs are yet to be trained. CNAPE could be improved by for instance incorporating allelic frequency information of common SNPs locating within these regions.

CNAPE takes a prior knowledge-aided multinomial logistic regression with LASSO to predict CNA, which differs from canonical DNA-seq based methods in that CNAPE first selects/inputs genes whose expression levels are responsive to copy number change, and then build regression models on these genes. The sparse-inducing LASSO filtered less informative genes, and prior-knowledge aided feature selection further improved the model performance, probably because it avoided selecting random artifacts and reduced the noise level. In future work, allele frequencies of SNPs will be included in the model to improve prediction accuracy, especially for smaller regions. The results from CNAPE would also be valuable to recalibrate tools such as GISTIC to identify regions with significant copy number aberrations from large cohorts [29–30].

## ACKNOWLEDGMENT

This work was supported by RGC grant No. 26102719, NSFC/RGC grant (No. N_HKUST606/17), CRF grants (No. C6002-17GF, No. C7065-18GF), the Hong Kong Epigenomics Project (EpiHK), and the Innovation and Technology Commission (ITCPD/17-9).

